# mRNA Imprinting: transcription apparatus can remotely control cytoplasmic post-transcriptional mechanisms by dozens of proteins

**DOI:** 10.64898/2026.02.03.703598

**Authors:** Shira Urim, Artyom Artamov, Shubham B. Deshmukh, Mordechai Choder

**Affiliations:** Department of Molecular Microbiology, Rappaport Faculty of Medicine, Technion-Israel Institute of Technology, Haifa, Israel

## Abstract

Proper regulation of gene expression requires the harmonious coordination of transcription with other stages of the mRNA lifecycle. We have previously demonstrated that proteins can bind to nascent transcripts co-transcriptionally – a phenomenon named “mRNA imprinting” because the bound proteins modulate subsequent stages of the mRNA lifecycle. Using a novel high-throughput approach, “PRofiling OF Imprinted Transcripts” (“PROFIT”), to identify proteins involved in mRNA imprinting, we uncovered several dozen candidates. The RNA polymerase II (Pol II) subunit Rpb4, which can itself imprint mRNAs, mediates the imprinting of a large subset of these proteins. Imprinting repertoire and profile is responsive to environmental changes and include HSP70 variants. Interestingly, PROFIT identified proteins previously thought to function mainly or exclusively in the cytoplasm, including a translation factors, mRNA decay factor, protein chaperones, substrate-delivering factors of the proteasome, targeting factors that deliver mRNAs to mitochondria and more. Using proximity labeling, we validated several hits, including two translation initiation factors: eIF4G and the eIF3 component Rpg1. Importantly, we found that the same Rpg1 molecule, which had been transiently localized near Pol II, co-sediments with polyribosomes - similar to the bulk Rpg1. Our results suggest that the transcription machinery can regulate translation by recruiting specific translation factors, which later participate in protein synthesis. mRNA imprinting appears to be a widespread phenomenon, and we speculate that it may not be limited to the transcription stage alone. Interestingly, the PROFIT experiments identified proteins with known **cytoplasmic** function, including a translation factor (Rpg1), mRNA decay factor (Xrn1), protein chaperones (Ssa1/2), substrate-delivering factors of the proteasome, targeting factors that deliver mRNAs to mitochondria, and actin-binding factors.

## Introduction

Effective coordination among the various stages of the RNA lifecycle is critical for proper gene expression. Co-transcriptional binding of factors with RNA polymerase II (Pol II) transcripts during transcription has been found to link transcription and post-transcriptional regulation of gene expression. This phenomenon is called “mRNA imprinting” (Choder, 2011). The Rpb4/7 heterodimer is a leading example of mRNA imprinting; it binds to nascent transcripts and accompanies them to impact post-transcriptional stages. Specifically, Rpb4/7 plays a role in transcription initiation, elongation, and polyadenylation (Choder, 2004; Orlicky et al., 2001; Pillai et al., 2001; Újvári and Luse, 2005) and is involved in the translation and decay of a particular class of transcripts following co-transcriptional RNA-binding (Goler-Baron et al., 2008; Harel-Sharvit et al., 2010; Lotan et al., 2007, 2005).

During the movement of mRNA/Rpb4/7 from one stage to the next, Rpb4/7 undergoes different post-translational modifications (Richard et al., 2021). We propose that these modifications serve as a communication language between the various stages of gene expression. Furthermore, we suggest that Rpb4/7, by participating in all stages of the mRNA lifecycle, can integrate them into a cohesive system (Richard et al., 2021). Other yeast proteins were found to bind Pol II transcripts co-transcriptionally, such as Ash1 (Shen et al., 2010) Recently, we reported that a “canonical” transcription factor (TF), Sfp1, imprints specific classes of mRNAs in a manner dependent on its binding to its canonical promoters. Unlike Rpb4/7, whose imprinting capacity is dependent on binding to Pol II, Sfp1 imprinting is dependent on its recruitment to the promoter, quite possibly through binding the transcription factor Rap1 (Kelbert et al., 2024). Recently, numerous DNA-binding TFs were shown to bind nascent RNA co-transcriptionally, thus affecting the nuclear stages of gene expression, notably chromatin occupancy (Oksuz et al., 2023). However, as of today, only a handful of factors were reported to bind mRNA co-transcriptionally and subsequently accompany and affect the mRNA lifecycle. Thus, the full scope of mRNA imprinting is unknown.

Also in higher eukaryotes, several cases have been reported of proteins that appear to bind Pol II transcripts co-transcriptionally while also functioning post-transcriptionally. For example, the principal targets of the mRNA-binding protein tristetraprolin (TTP) are pre-mRNAs rather than mature mRNAs. Since splicing typically occurs co-transcriptionally, it has been proposed that TTP binds its target mRNAs during transcription. Consistently, the nuclear TTP–pre-mRNA interaction is a prerequisite for the formation of TTP-containing mRNA complexes in the cytoplasm, where TTP regulates mRNA stability (Bestehorn, 2025). Similarly, FUS binds Pol II transcripts co-transcriptionally (Masuda et al., 2015), while TDP-43 functions in transcription and splicing ((Ratti and Buratti, 2016)). Both FUS and TDP-43 are also involved in various post-transcriptional stages, including cytoplasmic processes that the mRNP undergoes (Colombrita et al., 2012). Although, in all these cases, it remains unclear whether the same protein that regulates transcription subsequently dissociates from the transcription machinery to bind the RNA, this is a plausible scenario. It is also unknown whether the same protein that binds Pol II transcripts in the nucleus accompanies these mRNAs into the cytoplasm and functions there.

High-throughput approaches to identify interactomes of newly synthesized RNA - including nascent transcripts - have identified hundreds of RNA-associated proteins in mammalian cells (Bao et al., 2018; Heindel et al., 2023). However, these approaches could not distinguish between newly synthesized RNAs and those that remain physically associated with Pol II. In both cases, metabolic labeling, used to cross-link the labeled RNA with proteins, was performed for extended periods (16 hours), making the proportion of Pol II-bound RNA unclear and likely low. Shorter metabolic labeling times have been employed in PAR-CLIP–based analysis (Battaglia et al., 2017) or by purifying Pol II elongation complexes from a single gene locus (Harlen and Churchman, 2017). However, these approaches focused only on the known RNA biogenesis RNA proteins, (verifying that they bind nascent RNAs) (Battaglia et al., 2017), or to the analysis of transcription at a single locus (Harlen and Churchman, 2017).

Here, we introduce “PRofiling OF Imprinted Transcripts” (PROFIT) and BioPROFIT, a proteome-wide approaches that directly capture proteins bound to Pol II–associated nascent RNA. These methods enabling systematic and unbiased interrogation of the co-transcriptional RNA interactome.to identify yeast proteins that bind Pol II transcripts co-transcriptionally and later remain bound to the mature mRNAs. We found several such proteins that are known to regulate only transcription and were not considered as imprinting factors. We also found other proteins of known cytoplasmic functions. We found that many imprinting events are mediated by Rpb4 and are responsive to the environment. We focused on the transcription factor Spt6 and the translation factor Rpg1 as cases in point. We demonstrated that Spt6 imprinting capacity is compromised by a point mutation in Rpb4 that interfere with Spt6-Rpb4 interaction. We verified various PROFIT hits by proximity labeling (PL), including the translation factors Tif4631 (eIF4G) and Rpg1 (eIF3 subunit). Utilizing the PL method we addressed the question whether the interaction of Rpg1 with Pol II is at all related to its functions in translation. We thus discovered that the biotinylated Rpg1, which became biotinylated due to its proximity to Pol II, was associated with polyribosomes, much like the bulk Rpg1.

## Results

### Capturing of the newly transcribed RNA interactome by “PRofiling OF Imprinted Transcripts” (“PROFIT”)

In this study, we sought to identify the nascent RNA interactome of protein-encoding transcripts, which emerge from RNA polymerase II (Pol II). We developed a direct and unbiased approach to investigate nascent RNA-binding proteins.

To systematically identify the proteins that co-transcriptionally bind to nascent mRNAs, we developed an unbiased biochemical method, which we abbreviate as “PROFIT” (PRofiling OF Imprinted Transcripts), schematically illustrated in Fig. 1A and described in its legend and Materials and methods. Briefly, cells, carrying a Rpb3-FLAG-Tagged Pol II are cryogenically ground (Chattopadhyay et al., 2022). To stabilize protein-RNA interaction we performed a UV crosslinking (Urdaneta and Beckmann, 2020; Van Ende et al., 2020). Encouraged by the success of UV cross-linking of frozen tissue (Urdaneta et al., 2019) and in order to prevent any physiological response of the live cells to the irradiation, we performed the crosslinking on the frozen grindate (on dry ice). The grindate is then dissolved and the chromatin is fragmented by RNase-free DNase I digestion. FLAG-Tagged Pol II is isolated from the chromatin digest, carrying ∼ 50 bp DNA that is protected by Pol II (results not shown). Rnase I is then used to release the proteins bound to the transcriptional apparatus via the nascent transcript. The resulting proteins was analyzed using liquid chromatography-tandem mass spectrometry (LC-MS/MS) or western blot.

**Figure 1.**
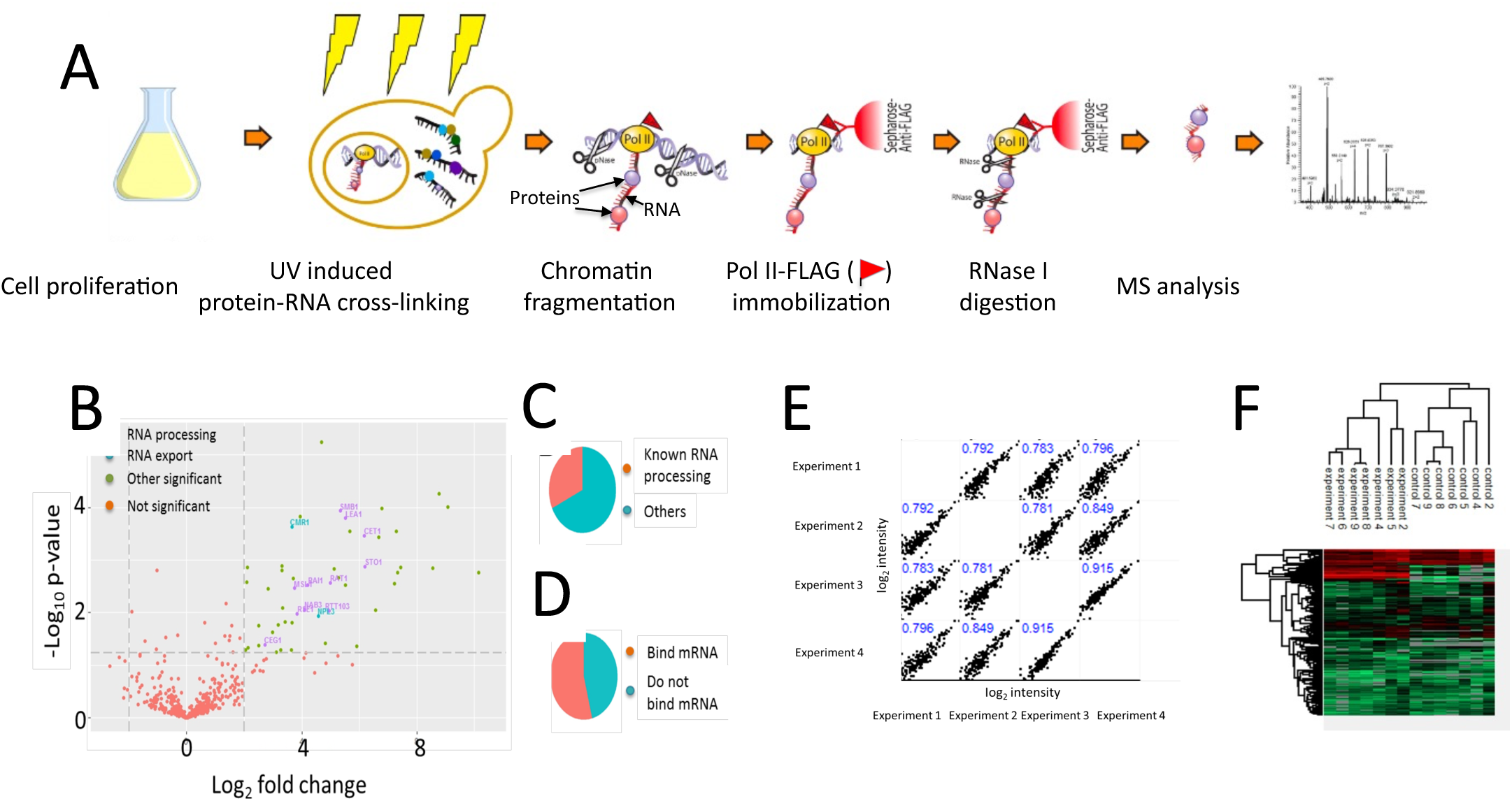
PROFIT identifies proteins known to involve in mRNA biogenesis and environmentally responsive factors with known cytoplasmic functions. **(A) *Schematic of the PROFIT approach***. Following UV crosslinking and cell lysis, chromatin was fragmented by RNase-free DNase treatment. The Pol II–DNA–RNP ternary complex was immobilized via the FLAG-tagged Rpb3 subunit using anti-FLAG antibodies. Nascent RNA–binding proteins were released by on-column RNase I digestion and identified by mass spectrometry (MS). **(B) *Volcano plot of significant PROFIT hits***. Proteins released by RNase I digestion of immobilized FLAG-tagged Pol II complexes were compared with a no-tag control strain. MS data from four biological replicates were analyzed using MaxQuant and Perseus. Statistical significance was assessed by a two-sample *t*-test with permutation-based FDR correction (FDR = 0.5, S0 = 0.1). Selected RNA-processing factors are indicated. **(C) *Distribution of significant hits classified as known RNA-processing factors (Battaglia et al., 2017) or other proteins.* (D) *Distribution of significant hits according to their reported ability to bind mature poly(A)+ mRNAs, based on published datasets (Beckmann et al., 2015; Mitchell et al., 2013a) and this study.* (E) *Reproducibility of RNase I–released protein intensities across biological replicates***. Pearson correlation coefficients are shown. **(F) *Hierarchical clustering and heat map of PROFIT LFQ intensities***. Experimental and control samples segregate into distinct clusters. LFQ data were analyzed using Perseus following MaxQuant processing. Proteins detected in fewer than three replicates in at least one group were excluded; missing values were imputed from a normal distribution.

Statistical analysis of at least four replicates is carried out using Perseus software and the results were expressed by using volcano plot (Fig. 1B). Encouragingly, we identified many proteins that were known to bind nascent mRNA transcripts during transcription (named “RNA Processing” in Fig. 1C), including capping enzymes, splicing factors, transcription elongation factors, polyadenylation factors and transcription termination factors (see Supplemental Table 2). Many of these proteins were identified by other investigators, either by using a PAR-CLIP analysis (Battaglia et al., 2017), or by purifying Pol II elongation complexes from a single gene (Harlen and Churchman, 2017). Unlike our unbiased approach, these published approaches focused only on the studied RNA biogenesis RNA proteins (verifying that they bind nascent RNAs) (Battaglia et al., 2017) or on a single gene (Harlen and Churchman, 2017).

Interestingly, the PROFIT experiments identified proteins with known cytoplasmic functions, including translation factor (Tef4631, Rpg1), mRNA decay factor (Xrn1), protein chaperones (Ssa1 and Ssa2), substrate-delivering factors of the proteasome, targeting factors that deliver mRNAs to mitochondria.

PROFIT analysis identified ∼40 co-transcriptionally associated proteins (Supplemental Table 2). These proteins include mRNA processing factors such as subunits of the mRNA capping enzyme, Cet1 and Ceg1, splicing factors such as Lea1 and Smb1, the quintessential imprinting factors Rpb4/7, and other nascent RNA processing factors (Figure 1C, Supplemental Table 2). The transcription elongation factor Spt6 was also identified. In addition to its role in the regulation of transcription, Spt6 has recently been implicated in mRNA turnover and was proposed to regulate mRNA buffering. Specifically, the absence of Spt6-Pol II interaction with Pol II CTD caused a decrease in the synthesis of a range of genes, whereas some of these mRNAs were more stable (Dronamraju et al., 2018).

In addition to RNA processing factors, several proteins with functions unrelated to transcription or RNA processing have been identified. For example, Xrn1p, Rat1p, and Rai1p, which are involved in RNA degradation, were identified. In addition, two translation initiation factors were identified. Notably, upon comparison of the ROFIT hits with published mature mRNA interactome datasets, considerable overlap was observed (Figure 1D).

### Proximity labeling reveals that several imprinting factors are proximal to Pol II

To obtain *in vivo* support for the PROFIT results and to determine which of the hits is proximally located to Pol II (perhaps transiently), we applied a proximity labelling (PL) method, which is based on the BirA-AviTag system. The WT form of BirA is a highly specific biotin ligase. It specifically biotinylates an *E. coli* substrate, 15-AA of which are enough for recognition and biotinylation. This AA sequence, named “AviTag”, can serve as a tag that becomes biotinylated by BirA, provided that the tag is within 10-15 nm of its enzyme (Cull et al., 2000).

We surgically fused BirA to the c-terminal disordered domain (CTD) of Rpb1, at *RPB1* locus (Fig. S1A), assuming that CTD flexibility will increase biotinylation radius and biotinylation efficiency. The CTD was found to accommodate liquid-liquid phase separated droplets at various stages of the transcription cycle (Li et al., 2024). We surmised that if the AviTagged protein is located in the same droplet, labeling would be efficient. This fusion had little or no effect on cell proliferation (Fig. S1B). To determine whether Rpb1-BirA can biotinylate PROFIT hits, we examined several hits, and observed that ∼65% of them were biotinylated during short labelling times, including a Hsp70 variants Ssa1 and Ssa2 (Fig. S1D), a transcription initiation factor Taf14, an mRNA decay factor Xrn1 (Figure S1E) and transcription elongation factor Spt6 (Fig. S1F). We note that although biotinylation of these PROFIT hits supports Pol II proximity, the lack of biotinylation does not rule out co-transcriptional binding; instead, it suggests that not all imprinting events involve a stage proximal enough to the BirA fused Pol II. Rpb1-BirA efficiently biotinylated most PROFIT hits, except for Rpg1-AviTag and Tif4631-AviTag. We found that Rpb9-BirA was more efficient in biotinylating the latter translation factors (Fig. S1C). Biotinylation of translation factors by Pol II subunit poses a specific issue. It is possible that it is not only the nuclear Pol II-BirA fusion that biotinylates these factors, but also the newly synthesized BirA fusion that naturally resides proximal of the translation apparatus. However, since BirA is fused to the C-terminal of the Pol II subunit, as soon as it is fully synthesized and properly folded, it left the translation apparatus. Nevertheless, to rule this possibility out, we determined biotinylation of Rpg1 after translation was blocked by cycloheximide (CHX). Indeed, no change in the extent of biotinylation was observed between CHX treated and untreated samples (Fig. S2). We conclude that Rpg1 biotinylation occures outside the context of translation.

### Proximity labeling validates imprinting and shows that Bio-Rpg1-Avi co-sediments with polysomes

In addition to the ability of PL to provide additional support for PROFIT, an advantage of PL lies in its capacity to track the fate of the protein after it leaves the transcription apparatus. In cases where proximity to Pol II is transient and the protein accompanies the mRNA post-transcriptionally—a criterion of mRNA imprinting—the biotin can serve as a history mark and help decipher the function of the imprinting protein (Fig. 2B). We followed the fate of Bio-Rpg1-Avi in the cytoplasm, as a case in point. Since Rpg1 is a component of eIF3, we examined the association of Bio-Rpg1-Avi, whose biotinylation was carried out by Rpb9-BirA, with polysomes. Cells, proliferated in biotin-depleted medium, were pulse-labeled with biotin for either 4, or 25 min, or continuousely labeled in biotin-rich medium (YPD) (Fig. 3). First, we determined the distribution of the bulk Rpg1-Avi using anti-AviTag antibodies and found, as expected (Li et al., 2009) that it mainly co-sedimented with the lighter fractions including the 40S, 60S ribosome subunits and the 80S monosome. We also observed strong association of Rpg1-Avi with the last heavier factions of the polysomes, which may be due to Rpg1 association with condensates (e.g., stress granules). We then examined Bio-Rpg1-Avi, using infrared fluorecent streptvidin (IR-streptavidin), and found that, like the bulk Rpg1-Avi, also Bio-Rpg1-Avi co-sedimented with ribosomes. Co-sedimentation was time dependent: at 4 min post-labeling, Bio-Rpg1-Avi was enriched in the light fractions of the gradient. Only after 25 min did the distribution of Bio-Rpg1-Avi become comparable to that of bulk Rpg1-Avi, detected by anti-AviTag antibodies (Fig. 3B–C). We calculated the ratio between bio-Rpg1-Avi and the bulk Rpg1-Avi for each fraction and found different patterns depending on the labeling time. During continuous labeling, the ratio was similar throughout the gradient, suggesting that imprinted Rpg1 associates with polysomes similarly to the bulk Rpg1. Following the shortest pulse labeling of 4 min, the ratios in the light fractions were higher than in the heavier fractions, consistent with the association of newly labeled Bio-Rpg1-Avi mainly with ribosome-free mRNAs. Longer labeling times shifted the pattern towards that of continuous labeling (Fig. 3C). These results are consistent with labeling of Bio-Rpg1-Avi in the nucleus, away from the translation apparatus, such that time is required for Bio-Rpg1-Avi to assemble with the translation machinery.

**Figure 2.**
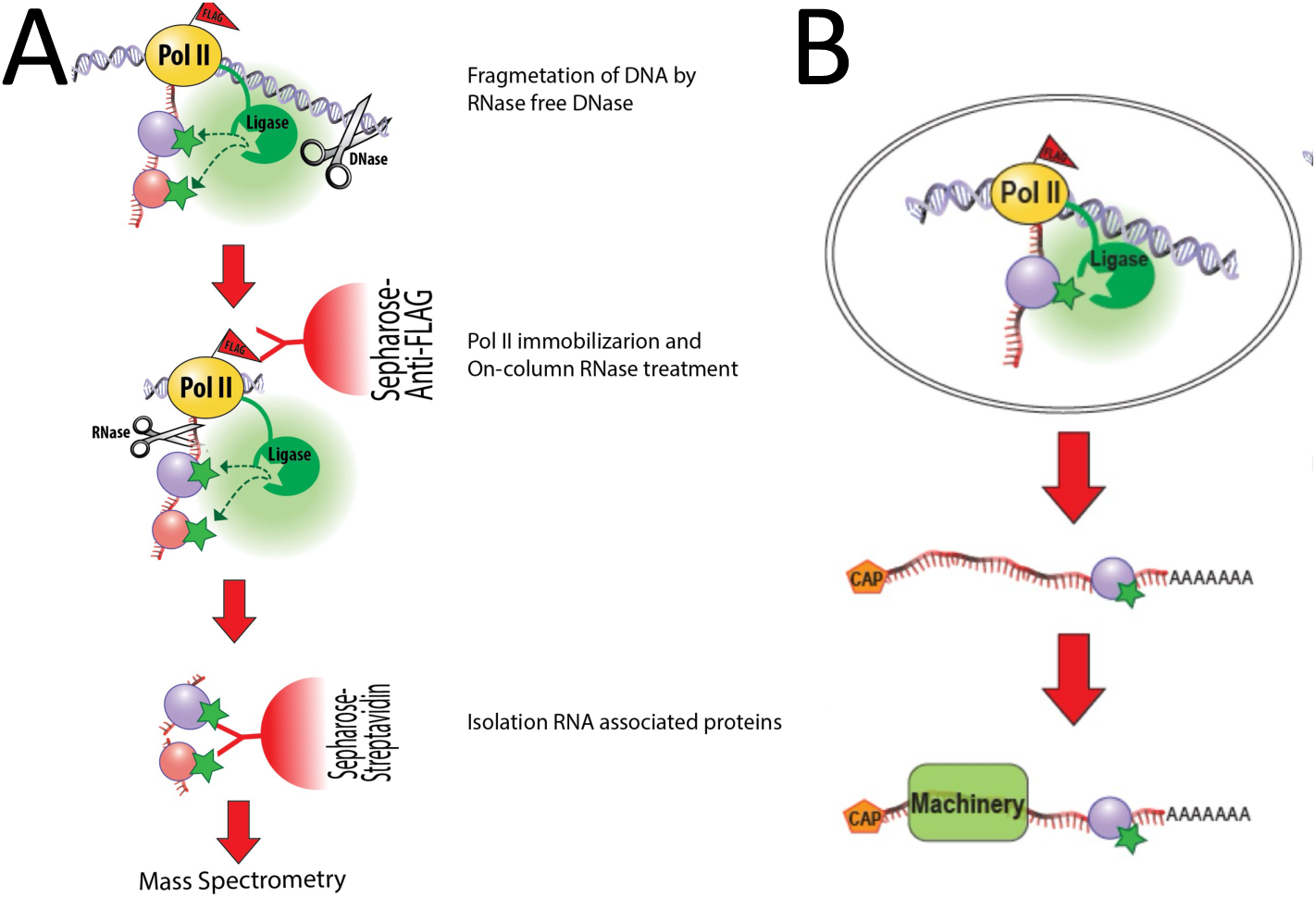
BioPROFIT combines PROFIT with proximity biotinylation to record transcription-proximal interactions and enables to follow the fate of the imprinting protein. **(A) Schematic of the BioPROFIT approach.** As in PROFIT (Fig. 1), Pol II complexes were purified via Rpb3-FLAG. In addition, BirA (codon-optimized for *S. cerevisiae*) was fused to the C-terminal domain (CTD) of the Pol II subunit Rpb1. BioPROFIT signal therefore depends on both in vivo proximity-dependent biotinylation and in vitro Pol II purification. **(B) *Biotin as a molecular “history mark”***. A PROFIT hit fused to eight AviTags is biotinylated when it resides in proximity to Rpb1-BirA during transcription. This design enables tracking of proteins that engage nascent transcripts and subsequently participate in post-transcriptional processes.

**Figure 3.**
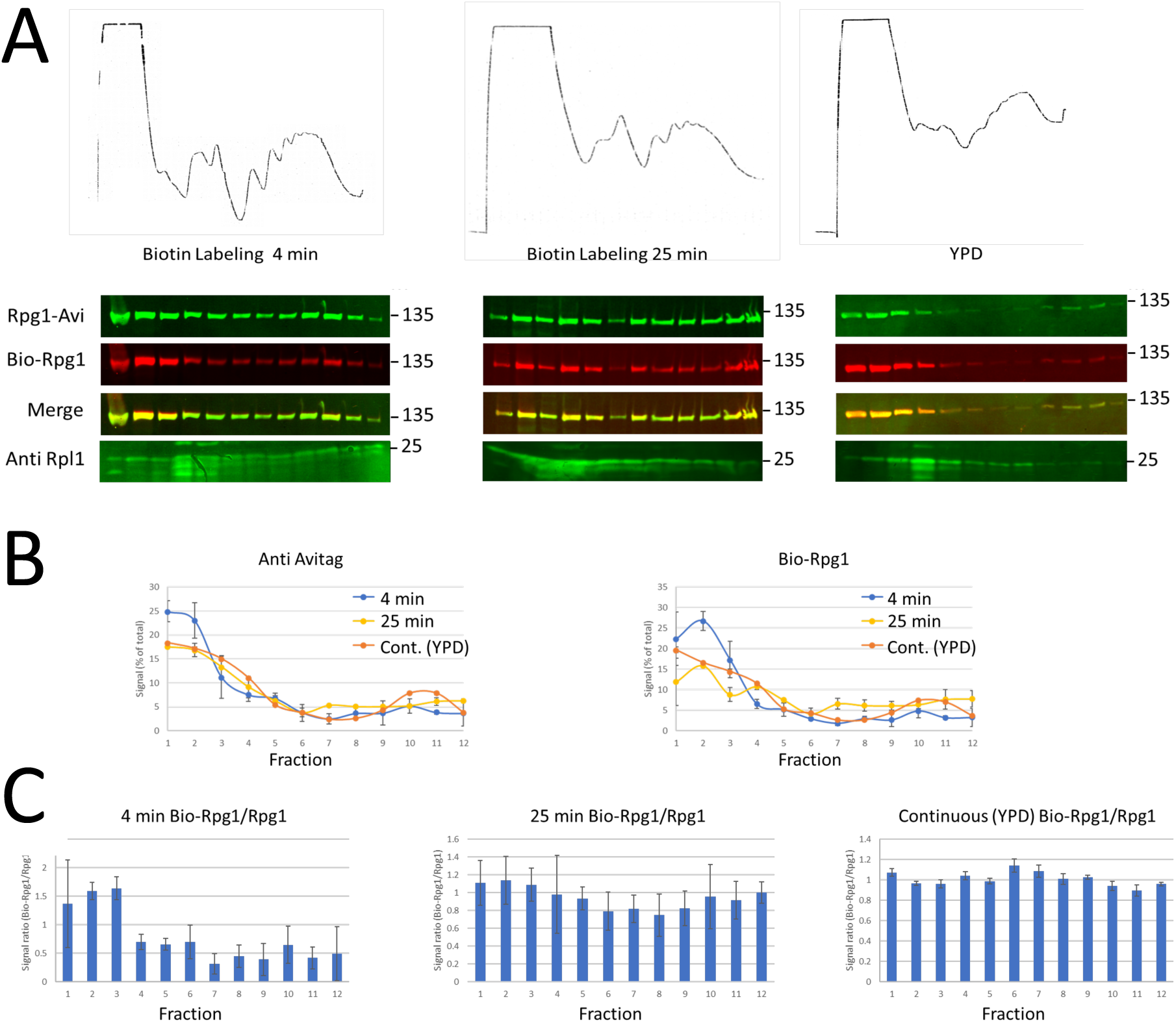
Rpg1 co-sediments with polysomes. *RPB9*-BirA, *RPG1*-AviTag cells were grown in minimal biotin medium and labeled with biotin for 4 or 25 min, or continuously grown in YPD medium containing excess biotin. Cell extracts were subjected to sucrose-gradient polysome fractionation as described in *Materials and Methods*. **(A) *Polysome fractionation.*** Upper panels: polysome profiles. Lower panels: gradient fractions were subjected to western blot assay. Membranes were probed with IR-streptavidin (Bio-Rpg1-Avi), α-AviTag (bulk Rpg1-Avi), and α-Rpl1 antibodies (ribosomal protein L1). The position of the molecular weight marker is indicated. **(B) *Gradient profiles of Bio-Rpg1-Avi (right panel) and bulk Rpg1-Avi (left panel).*** Band intensities were quantified (*Materials and Methods*) and normalized such that each fraction represents a percentage of the total signal across the gradient. **(C) *Comparison of Bio-Rpg1-Avi and bulk Rpg1-Avi sedimentation.*** Profiles differ after 4 min of biotin labeling but converge at later time points. Ratios of normalized Bio-Rpg1-Avi to normalized bulk Rpg1-Avi were calculated for each fraction; a value of 1 indicates identical sedimentation.

Co-sedimentation of Bio-Rpg1-Avi with ribosomes was sensitive to the translation inhibitor puromycin, which stimulates the release of the nascent peptide followed by ribosome dissociation (Blobel et al., 1971). As shown in Fig. 4, puromycin treatment led to an increase in free subunits and a shift in the sedimentation of the ribosomal protein Rpl1 and the poly(A)-binding protein Pab1 to the lighter fractions. Both the bulk Rpg1-Avi, detected by anti-AviTag Abs and the Bio-Rpg1-Avi, detected by IR-sreptavidin similarly shifted to the lighter fraction, corroborating the association of Bio-Rpg1-Avi with ribosomes. Puromycin sensitivity is consistent with association of Bio-Rpg1-Avi with translationally active polysomes. Collectively, these results are consistent with co-transcriptional binding of Rpg1-Avi with mRNA that later is engaged in the translation apparatus outside the nucleus, indicating that Rpg1-Avi can imprint mRNAs.

**Figure 4.**
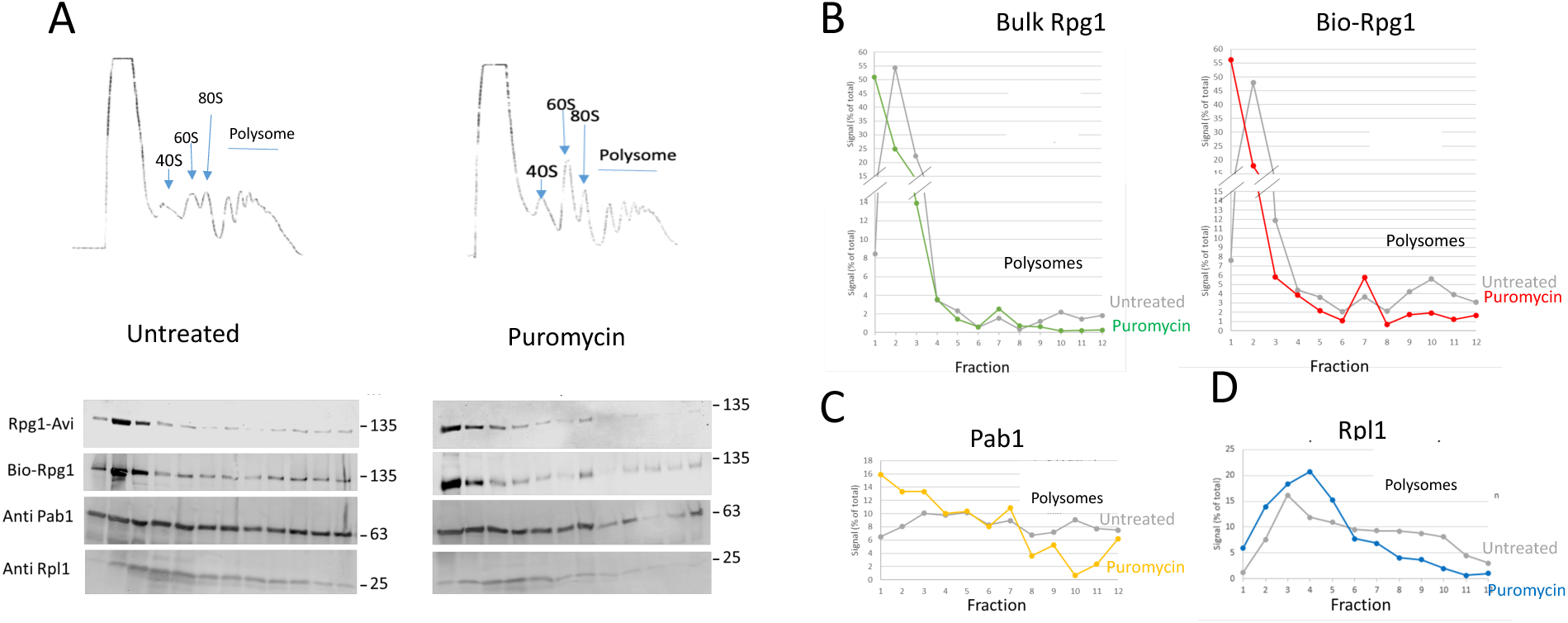
Bio-Rpg1-AviTag co-sediments with polysomes in a puromycin-sensitive manner. **(A)** Rpb9-BirA, Rpg1-AviTag cells were grown in minimal biotin medium and labeled with biotin for 10 min prior to harvest. Cell lysates were divided into untreated and puromycin-treated (1.6 mM, 15 min) samples and subjected to sucrose-gradient polysome fractionation. Upper panels: polysome profiles. Lower panels: gradient fractions were subjected to western blot assay. Membranes were probed with IR-streptavidin (Bio-Rpg1-Avi), α-AviTag (bulk Rpg1-Avi), α-Rpl1 (ribosomal protein L1), and α-Pab1 antibodies. The position of the molecular weight marker is indicated. **(B–D) Quantification of gradient fractions.** Band intensities were normalized to the total signal per gradient and are presented as percentages.

### Co-transcriptional binding of proteins to Pol II transcripts is responsive to the environment

To determine whether the profile of PROFIT hits is responsive to the environment, we challenged cells with heat shock (HS). Co-transcriptional nascent RNA binding of most PROFIT hits decreased in response to HS, either because global transcription was downregulated (Choder and Young, 1993; Gasch et al., 2000)) or due to additional regulatory mechanisms. In contrast, co-transcriptional nascent RNA binding of a few proteins increased in response to the stress (Fig. 5A). Two Hsp70 variants, Ssa1 and Ssa2, showed increased nascent RNA binding, despite little increase in protein level (Fig. 5B input, and results not shown). In addition, several other factors were found to specifically associate with nascent transcripts during HS. These factors include the mitochondrial proteins Por1, as well as Adk1, and Egd2, which direct nascent peptides to the ER (Fig. 5 supplemental Table 3).

**Figure 5.**
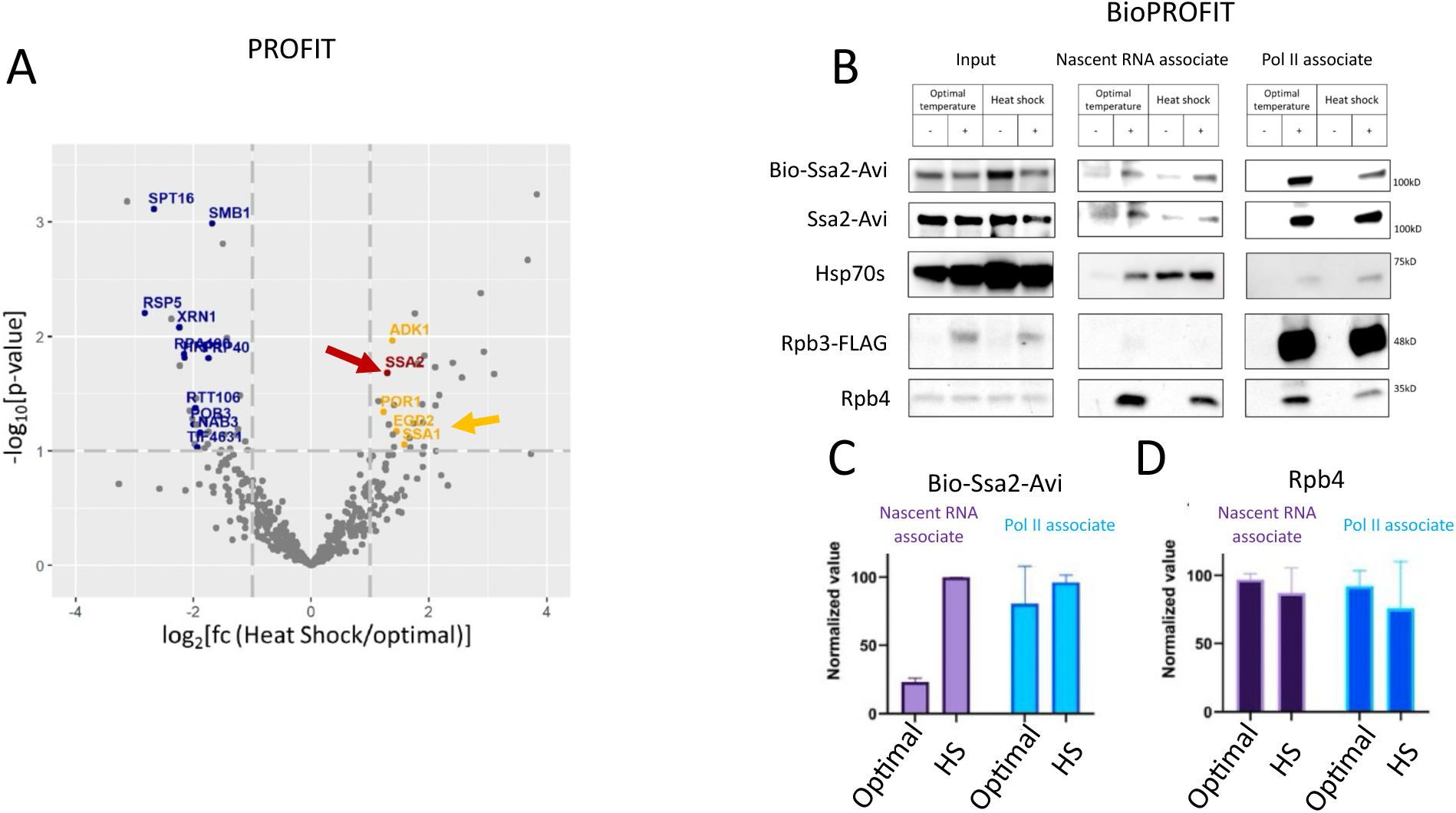
PROFIT and BioPROFIT reveal heat-shock-induced co-transcriptional binding of Ssa2 to nascent RNA. **(A) PROFIT analysis of heat-stressed versus non-stressed cells.** Cells were grown to mid-log phase and either harvested directly or shifted to 40 °C for 30 min prior to harvest. Cell grindates were subjected to PROFIT analysis (see *Materials and Methods* and Fig. 1A–B). Arrows indicate Ssa1 (yellow) and Ssa2 (red). **(B) *BioPROFIT identification of biotinylated Ssa2 (Bio-Ssa2) associated with Pol II (right) or nascent RNA (middle)***. Cells were grown in minimal biotin medium at the optimal temperature and either harvested directly or subjected to heat shock (40 °C, 30 min). Biotin (1 µM) was added for 15 min before harvest. Cells were processed as in Fig. 1A. Following RNase treatment and analysis of the release proteins (“Nascent RNA associated”), the RNP-depleted Pol II complex, remained bound to the column, was eluted with FLAG peptide (“Pol II–associated”). Proteins were analyzed by western blot. The same membrane was decorated with IR-streptavidin to detect Bio-Ssa2-Avi; with anti-Hsp70 antibodies to detect Ssa2-Avi and other Hsp70s (distinguished by their electrophoretic mobility); anti-FLAG antibodies to detect Rpb3-FLAG; and with anti-Rpb4 antibodies to detect Rpb4. **(C) *Quantification of Bio-Ssa2 associated with nascent RNA and Pol II***. Bio-Ssa2 was detected using IR-streptavidin, and Rpb3-FLAG was detected using anti-FLAG antibodies. Bio-Ssa2 values were normalized to the corresponding Rpb3-FLAG signal at each temperature. For each sample pair, the maximal value was set to 100%. **(D) *Quantification of Rpb4 associated with nascent RNA and Pol II, normalized to Rpb3-FLAG as in (C)*.**

To verify co-transcriptional binding of Ssa2 to nascent RNA by independent means, we tagged *SSA2* with AviTag and performed BioPROFIT, which combines proximity labeling with PROFIT (Fig. 2), two independent processes. Figure 5B (“Pol II–associated” panel) shows that interaction of Ssa2-Avi with Pol II is detected by both approaches - affinity purification and proximity labeling. Moreover, Bio-Ssa2-Avi can be released from the Pol II ternary complex by RNase digestion (Fig. 5B, “Nascent RNA–associated” panel). Quantification indicates that binding to nascent RNA, relative to Pol II levels, increases during HS (Fig. 5C). This increase is mainly due to a decrease in chromatin-bound Pol II, consistent with general transcriptional repression in response to HS (Choder and Young, 1993; Gasch et al., 2000). Notably, in response to HS, Bio-Ssa2-Avi binding to Pol II decreased, whereas binding to nascent RNA did not, suggesting a relative shift of Ssa2 association from Pol II toward nascent RNA during HS. In contrast to Ssa2, nascent RNA binding of Rpb4, whose imprinting was demonstrated previously (Goler-Baron et al., 2008; Harel-Sharvit et al., 2010; Lotan et al., 2007, 2005), did not change during HS; the apparent decrease in nascent RNA binding paralleled the decrease in Pol II levels. Results obtained by decorating the membrane with anti-Hsp70 antibodies show that, within the limits of western blot assay, less untagged Hsp70, migrating as a 70 kDa band, was detected compared with Ssa2-Avi, which migrated as a ∼105 kDa band (Fig. 5B, cf Hsp70 band with Ssa2-Avi). We suspect that Ssa3 and Ssa4, which are substantially induced during HS, do not bind nascent RNA for the following reasons: (1) they were not identified in our PROFIT approach, whereas Ssa1 was and (2) the intensity of the Pol II-bound Hsp70 band did not increase during HS. We therefore suspect that the 70kD band is composed mainly of Ssa1.

### PROFIT identifies Rpb4 as a central mediator of co-transcriptional binding to nascent RNA

Approximately half of the proteins associated with nascent RNA have been found to interact with Rpb4 (Supplemental Table 4). As the nascent RNA exits from Pol II active site, it interacts with Rpb4/7 (Chen et al., 2009; Újvári and Luse, 2005), placing Rpb4/7 in a good position to mediate mRNA imprinting. This raised a possible role for Rpb4/7 in regulating mRNA imprinting and prompted us to compare PROFIT in cells carrying either a deletion or mutations in *RPB4*. Indeed, we found many factors that were differentially imprinted (Fig. 6A and Supplemental Table 4). Remarkably, most of them are known Rpb4-interacting proteins (Supplemental Table 4). The implications of this finding is discussed in the Discussion.

**Figure 6.**
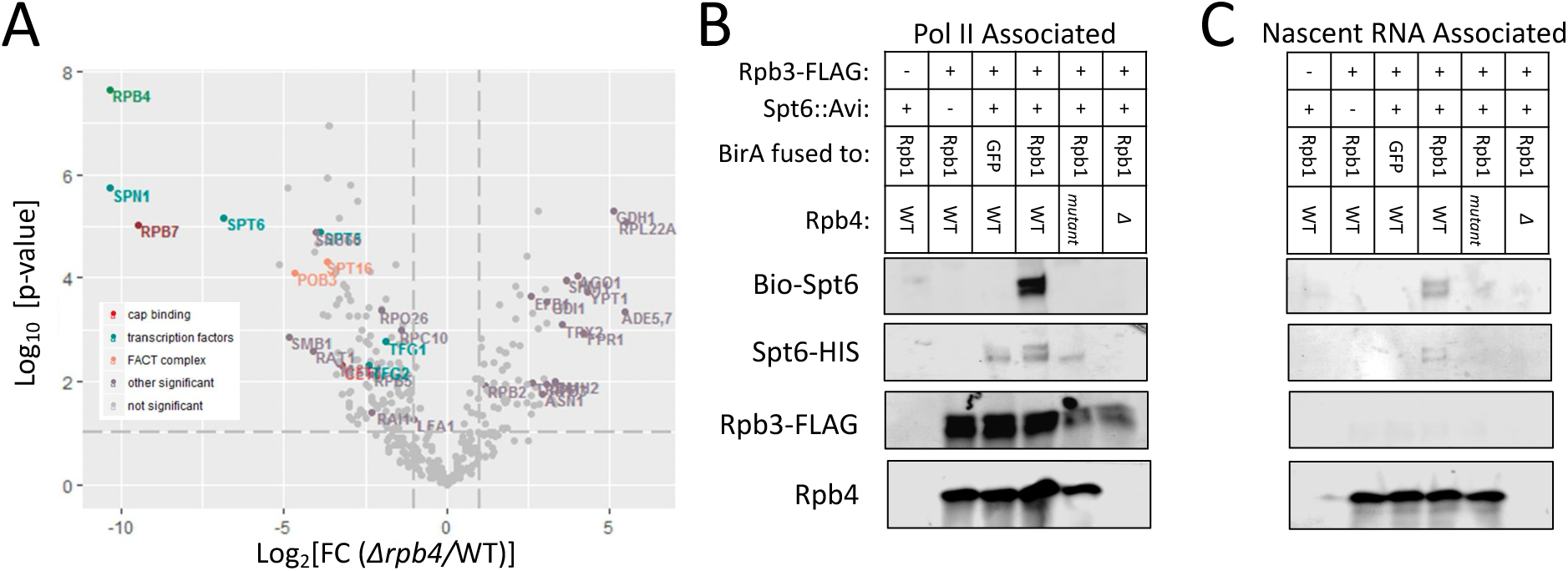
Rpb4 is required for co-transcriptional binding of multiple PROFIT hits. **(A) *Volcano plot comparing PROFIT results from WT and rpb4Δ cells***. Grindates of *RPB3*-FLAG cells carrying or lacking *RPB4* were subjected to PROFIT analysis as described in Fig. 1A and in *Materials and Methods*. Statistical analysis of two biological replicates (each including experimental and no-tag controls) was performed using Perseus. **(B) *BioPROFIT analysis showing that efficient co-transcriptional binding of Spt6-Avi to nascent RNA depends on Rpb4***. Cells, genotypes of which are indicated (“mutant” refers to or *rpb4*^E19D, E20D, E21D, E22D^), were grown in minimal biotin medium under optimal conditions. Biotin (1 µM) was added for 15 min before harvest. Cells were processed as in Fig. 1A. Following RNase treatment and analysis of the release proteins (“Nascent RNA associated”), the RNP-depleted Pol II complex, remained bound to the column, was eluted with FLAG peptide (“Pol II–associated”). Proteins were analyzed by western blot. The same membrane was decorated with IR-streptavidin to detect Bio-Spt6-Avi; anti-FLAG antibodies to detect Rpb3-FLAG; and with anti-Rpb4 antibodies to detect Rpb4. The bulk Spt6 was detected by anti-HIS Abs that bound to (HIS)x6, which was placed upstream of the FLAG.

Among the factors whose association with the nascent transcript was decreased in *rpb4Δ* cells was Spt6. Spt6 interacts with Pol II, adjacent to the Rpb4/7 heterodimer (Vos et al., 2018). Given that the absence of *RPB4* resulted in a 7-fold reduction in Spt6 co-transcriptional association with nascent RNA determined by PROFIT (p= 2.14 × 10^−5^ see Supplemental Table 4) and that it was reported to bind mRNAs and affect post-transcriptional stages (Dronamraju et al., 2018), we examined Spt6 as a case in point for a factor whose imprinting is dependent on Rpb4, using BioPROFIT.

First, we created a strain expressing *RPB1*-BirA, *RPB3*-FLAG and *SPT6*-AviTag. We affinity purified Rpb3-FLAG-containing Pol II and found that the bulk Spt6-Avi, detected by anti-AviTag, was co-purified, as expected (Dronamraju et al., 2018; Sun et al., 2010), in a FLAG-dependent manner. It migrated as two bands, probably reflecting different post-translational modifications, as was reported previously (Dronamraju et al., 2018). Reacting the membrane with IR-streptavidin revealed that Bio-Spt6-Avi was co-purified as well (Fig. 6, “Pol II associated” panel). Its biotinylation was dependent on Rpb1-BirA and on the AviTag. In the absence of *RPB4*, Spt6-Avi was no longer co-purified with Pol II, indicating that Rpb4 is a critical component for the interaction of Spt6 with Pol II.

Previously, we reported the presence of an unstructured loop in Rpb4, carrying E19, E20, E21, E22 (“E19-22”) motif, mutations in which compromised interactions with a number of factors, including Spt6 (Richard et al., 2021). Unlike *rpb4Δ*, transcription in this mutant is not slower than in WT cells (Richard et al., 2021). Less Spt6-Avi was co-purified with this mutant Pol II (Fig. 5B, lane designated “mutant”), consistent with Rpb4 stabilizing Spt6-Pol II interaction. Interestingly, this small amount of Spt6-Avi was not biotinylated with Rpb1-BirA, suggesting that the interaction of Spt6-Avi with this mutant Pol II occurred at an abnormal position – far away from the BirA.

To determine whether the bio-Stp6 can bind nascent RNA co-transcriptionally and whether Rpb4 plays a role in this possible binding, we digested, in column, the affinity purified Pol II-containing complex with Rnase I. Bio-Spt6-Avi was released from the WT Pol II-containing complex, provided that its Rpb1 was fused to BirA. However, no bio-Spt6-Avi was released from the complexes containing Pol II mutants (Fig. 5C). Taken together, both PROFIT and PL lead to the same conclusion: Spt6-Avi interactions with both Pol II and nascent RNA are mediated by Rpb4.

## Discussion

### PROFIT revealed known and unexpected hits

RNA imprinting is a relatively new concept describing cross-talk between the transcription machinery and post-transcriptional events in the mRNA lifecycle. Our PROFIT method revealed several dozens hits. PROFIT identified both proteins that bind nascent RNA transiently -without remaining associated after its maturation - and proteins that remain bound to mature mRNAs, potentially affecting transcript fate. Examples of the former type are Spt16 and Pob3, which constitute the FACT complex, function as histone chaperones facilitating transcription (Zhou et al., 2020). To our knowledge, this is the first indication that FACT associates with nascent RNA, as the current literature provides no evidence that FACT directly binds RNA. We speculate that this nascent RNA-binding feature may help tether the FACT complex, while complexed with histones, near the elongating Pol II, thereby facilitating nucleosome reassembly behind Pol II. This may be analogous to the recruitment of regulatory factors by lncRNAs, which guide them to specific chromatin locations (Davidovich and Cech, 2015). Other examples include splicing factors (Smb1, Lea1), transcription elongation factors (Spt5 and Spt6), and transcription termination factors (Rtt103) (see Supplemental Table 2)

The two subunits of the capping enzyme (Cet1 and Sto1), which cap the nascent RNA during early transcription (Kim et al., 2004), represent additional examples. Whereas they bind the 5′ end of the nascent RNA for capping, there are no indications in the literature that these proteins bind mature yeast mRNAs. Instead the 5’ cap binds other factors that accompany the mRNA, like CBC and eIF4E; **t**herefore, we do not regard their transient binding as imprinting. We note, however, that according to a study posted on bioRxiv, the mammalian capping enzyme is a shuttling factor that functions both in the nucleus and the cytoplasm, where it can maintain capping homeostasis (Gayen et al., 2024). This raises the possibility that, in mammals, the nuclear enzyme that binds transcripts co-transcriptionally may imprint them.

We surmise that, for any given gene, co-transcriptional binding to nascent Pol II transcripts varies in efficiency. Imprinting by Xrn1, one of the PROFIT hits, was found to range between 0 and 100% (Chattopadhyay et al., 2022), and we suspect that this represents the norm for other imprinting factors. Indeed, imprinting appears to be regulated, as many PROFIT hits respond to environmental conditions (Fig. 5). This responsiveness suggests that imprinting plays a role in cellular adaptation to environmental changes. Notably, our approach is not ideally suited for detecting low-abundance proteins or proteins that imprint only a limited subset of mRNAs, due to signal-to-noise limitations (see Limitations of PROFIT). We therefore propose that mRNA imprinting is even more widespread than our current dataset suggests.

Many PROFIT hits were found to bind poly(A)+ mature mRNAs (see Supplemental Table 2). Examples include:

1. **Xrn1:** The major 5′–3′ exonuclease that degrades mRNA in the cytoplasm (Parker, 2012; Pérez-Ortín et al., 2013). We previously showed that Xrn1 also functions in transcription (Chattopadhyay et al., 2022; Fischer et al., 2020; Haimovich et al., 2013; Medina et al., 2014), binds ∼50% of nuclear nascent RNAs, accompanying them to the cytoplasm, and degrading them following decapping (Chattopadhyay et al., 2022). Preventing Xrn1 nuclear import substantially inhibits degradation of these mRNAs (Chattopadhyay et al., 2022), representing a classical case of mRNA imprinting. Xrn1 was found to bind nascent RNA by purifying Pol II elongation complexes from a single gene locus (Harlen and Churchman, 2017).
2. **Npl3:** Npl3 has been implicated in multiple processes of gene expression, including mRNA transcription elongation and termination (Bucheli et al., 2007; Dermody et al., 2008; Holmes et al., 2015) mRNA export under stress (Shen et al., 2000; Zander et al., 2016) and translation (Baierlein et al., 2013; Estrella et al., 2009). Npl3 is an RNA-binding protein that binds RNA in the nucleus, possibly co-transcriptionally (Keil et al., 2023; Moursy et al., 2023) and many of its known functions are carried out in the context of the mRNA. Importantly, Npl3 also binds Pol II by interacting with Rpb1 CTD (Gupta et al., 2023) and Rpb4 (Lotan et al., 2005). Together with our observation that it binds nascent RNA in Rpb4-dependent manner, these features support a classical imprinting model in which Npl3 accompanies Pol II during transcription elongation, transfers from Pol II to the emerging RNA, remains bound to the mRNA, and coordinates multiple post-transcriptional stages of the mRNA life cycle.

### Unexpected Examples

1. **Spt6:** While Spt6’s role in transcription elongation is established (Miller et al., 2023), it was recently implicated in mRNA turnover and buffering (Dronamraju et al., 2018). Spt6 binds Pol II, adjacent to Rpb1 CTD (Close et al., 2011) and Rpb4 (Vos et al., 2018). We found that Spt6-Pol II interaction is stabilized by Rpb4 (Fig. 6B). Spt6 recruits the Ccr4-NOT complex (Dronamraju et al., 2018), yet PROFIT did not identify Ccr4-NOT components, suggesting Spt6 may recruit Ccr4-NOT complex after mRNA maturation. We propose that Spt6 imprints target mRNAs, accompanies the target mRNA till they demise (Dronamraju et al., 2018), while influencing their stability through an unknown mechanism critical for mRNA buffering. Our model posits that imprinting efficiency is influenced by the transcription features. Thus, reduced transcription can lead to decreased Spt6 imprinting, and consequently increased stability of the imprinted transcripts. Indeed, cells expressing an Spt6 mutant, interaction of which with Pol II CTD is defective, show reduced synthesis of many genes; whereas the corresponding mRNAs exhibit increased stability and abundance (Dronamraju et al., 2018). This feedback mechanism should result in a balance between synthesis and decay reported by us(Blasco-Moreno et al., 2019; Chattopadhyay et al., 2022; Fischer et al., 2020; Goler-Baron et al., 2008; Haimovich et al., 2013; Kelbert et al., 2024; Lotan et al., 2005; Shalem et al., 2011) and others (Hartenian and Glaunsinger, 2019; Sorenson and Sieburth, 2025; Sun et al., 2012; Timmers and Tora, 2018). Rpb4 is essential for Spt6-mediated imprinting (Fig. 6), reinforcing the role of Rpb4 in this mRNA buffering (Shalem et al., 2011).
2. **Tif4631 Cdc33 (eIF4E) and Rpg1:** *TIF4631* and *CDC33* encode eIF4G and eIF4E respectively, subunits of the eIF4F complex that is critical for cap-dependent translation initiation (Das and Das, 2016). Rpg1 is a subunit of eIF3, also plays a key role in translation (Valášek et al., 2017). Our results suggest that the imprinting Rpg1 (Bio-Rpg1) functions similarly to bulk, mainly cytoplasmic, Rpg1 (Figs. 3 and 4). We propose that imprinting of Rpg1 mediates transcription–translation cross-talk: it binds mRNA co-transcriptionally and accompanies it until ribosome binding. Classical translation models posit the mRNA is recruited to the 48S ribosomal subunit post-eIF3 assembly (Valášek et al., 2017). Our model suggests otherwise for the imprinted mRNA, as it is already bound to Rpg1 during synthesis. eIF3 exists in a dynamic equilibrium of full complexes, subcomplexes, and individual subunits. Within the context of the full complement of eIF3, Rpg1 is necessary and sufficient for binding mRNA (Ide et al., 2024). Indeed, Rpg1 is the only eIF3 subunit that binds mRNA with high enough affinity in the cell, at least among its free subunits (Ide et al., 2024). Equally relevant to our work was the observation that Rpg1 stabilizes the interaction of the translated mRNA at both entry and exit channels of the 48S ribosomal subunit (Aitken et al., 2016), thus it seems to be sufficient for stabilizing the mRNA/small ribosomal subunit stabilization even in the absence of other eIF3 subunits These results and ours imply that imprinted Rpg1 nucleates eIF3 assembly around the mRNA following the export of mRNA/Rpg1 to the cytoplasm, either before or during its binding to the small ribosome subunit. Further research is needed: Do translation factors other than Rpg1, EiF4e and eIF4G bind Pol II transcripts co-transcriptionally, which escaped our detection? Is Rpg1 imprinting regulated? Is it transcript-specific? What are the mechanistic differences between standard translation, occurs exclusively in the cytoplasm, and imprinting-mediated translation?
3. **Rat1.** Rat1 resides mainly in the nucleus and can degrade RNA from 5’ to 3’, like Xrn1. Rat1’s co-transcriptional binding supports the “torpedo model,” where it degrades, from 5’ to 3’, the 3’ RNA fragment post-cleavage of the cleavage and polyadenylation stage (Luo et al., 2006). Rat1 has also a splicing role in some yeast genes (Dhoondia et al., 2021). Our ability to release Rat1 from Pol II by RNase is consistent with these two functions. Interestingly, Rat1 also binds mature poly(A)+ cytoplasmic mRNA (Mitchell et al., 2013; Shchepachev et al., 2019), raising the possibility that a portion of Rat1, possibly a portion that does not degrade the 3’ fragment of the cleaved RNA, binds the 5’ fragment and accompanies the RNA into the cytoplasm, akin to Xrn1 (Chattopadhyay et al., 2022). Rat1’s ability to replace Xrn1 under certain conditions (Johnson, 1997) supports this hypothesis.
4. Pol II subunits. Unexpectedly, PROFIT identifies nearly all Pol II subunits. A similar result was reported upon purification of Pol II elongation complexes from a single gene locus (Harlen and Churchman, 2017). The apparent association of Pol II subunits with mature mRNA, detected by high-throughput approaches (Supplemental Table 2), was equally unexpected. We suspect that this reflects an unrelated phenomenon, such as RNA-containing assembly intermediates. Alternatively, it may represent Pol II–mediated imprinting, as free Pol II complexes in the nucleoplasm could, in principle, bind the nascent RNA via intrinsic RNA-binding domains—for example, the Rpb7 RNA-binding domain (Choder, 2004).

### Rpb4/7 as a possible regulator of mRNA imprinting

The Rpb4/7 heterodimer occupies a strategic position within the RNA polymerase II (Pol II) holoenzyme. It is located near the Pol II C-terminal domain (CTD) and binds the nascent RNA as it emerges from the active site (Chen et al., 2009; Újvári and Luse, 2005), at least in some Pol II complexes. The effect of Rpb4 mutation on post-transcriptional regulation of the mRNA lifecycle is complex. On one hand, Rpb4/7 accompanies mRNA throughout its lifecycle, modulating its functionality by interacting with key regulatory proteins such as the translation factor eIF3 and the mRNA decay factors Pat1 and Lsm2 (Harel-Sharvit et al., 2010; Lotan et al., 2005). These activities appear to be regulated by various combinations of post-translational modifications (PTMs) that Rpb4/7 undergoes (Richard et al., 2021).

Here, we show that Rpb4 not only binds Pol II transcripts co-transcriptionally but also influences the mRNA lifecycle by mediating imprinting of other proteins. Strikingly, most PROFIT hits - imprinting of which depends on Rpb4 - are known Rpb4-interacting proteins (Supplemental Table 4). This suggests that their interaction with Rpb4 positions them near the emerging transcripts, giving them an advantage over other nuclear RNA-binding proteins. It is also possible that, following the dissociation of Rpb4/7 from the Pol II complex(Choder and Young, 1993; Goler-Baron et al., 2008; Harel-Sharvit et al., 2010; Selitrennik et al., 2006 Shalem), Rpb4 and some of these interacting PROFIT hits bind the emerging transcript cooperatively.

### Limitations of PROFIT

PROFIT is optimized to identify proteins that bind a sufficiently large fraction of nascent Pol II transcripts to surpass the method’s detection sensitivity. Consequently, proteins that associate with only a restricted subset of nascent transcripts—despite being biologically relevant—may fall below the statistical detection threshold and remain “under the radar.” An example is Sfp1, previously demonstrated to imprint ∼ 260 mRNAs (Kelbert et al., 2024), was detected in our mass spectrometric data but did not pass the stringent statistical cutoff applied in the PROFIT analysis. Likewise Ccr4-NOT complex (Reese, 2012; Trcek et al., 2011) She2 (Shen et al., 2010) and Dbp2 (Trcek et al., 2011)were not identified by PROFIT.

### Concluding remarks

In prokaryotes, transcription and translation are coupled through physical interactions (Blaha and Wade, 2025). In eukaryotes, however, these two processes occur in separate compartments. Despite this separation, eukaryotes appear to have retained cross-talk between these key mechanisms, albeit through a different evolutionary strategy. Recently, it was reported that distinct sets of newly transcribed mRNAs escape HS-induced translation repression, by an unknown mechanism (Glauninger et al., 2025). How cells identify newly transcribed mRNAs? We hypothesize that mRNA imprinting provides a plausible mechanism for tagging newly transcribed mRNAs and prevents their translation inhibition. Imprinting of translation factors serve as an attractive option.

We hypothesize that there is no *a priori* reason to assume that transcription is the only stage in the mRNA lifecycle that imprints the mRNA. In principle, imprinting might occur at any of the major stages. Specifically, it is possible that, during mRNP export through the nuclear pore complex, the mRNP binds (a) factor(s) that accompanies the mRNA and affects its fate. The same is true for any stage the mRNP undergoes. This expanded hypothetical view underscores a potential novel mechanism representing a flexible, modular regulatory strategy in gene expression.

## Acknowledgement

We thank all members of the Choder laboratory for fruitful discussions. We thank the Technion Genomics and Proteomics Centers for data generation, initial analyses, and technical advice. This work was supported by Israel Science Foundation, grant **310/20**.

## Materials and Methods

### Yeast strain construction

Yeast strains are listed in Supplemental Table 1. BirA was inserted at the 3′ end of the endogenous *RPB1* or *RPB9* loci without altering flanking non-coding regions. A RPB1::BirA::RPB1 3′NCR::URA3::RPB1 3′NCR cassette was integrated downstream of the final codon by homologous recombination and selected on uracil-deficient medium. URA3 was subsequently excised by recombination between identical 3′NCRs and selected on 5-fluoroorotic acid plates. Correct integration was confirmed by Sanger sequencing. AviTag was inserted by a similar approach.

### Yeast transformation

Yeast were transformed using the lithium acetate method as described previously (Choder and Young, 1993).

### Cell growth, harvesting, and UV crosslinking

Yeast cultures (3 L) were grown in YPD at 30°C with shaking (200 rpm) to mid-log phase. Cells were harvested by vacuum filtration onto 0.45-µm nitrocellulose membranes (Whatman), scraped, flash-frozen in liquid nitrogen, and cryogenically pulverized (six cycles, 30 s, 15 Hz) using a Retsch MM301 mixer mill. Pulverized material (“grindate”) was spread on liquid-nitrogen–chilled glass plates, placed on top of dry ice, and UV-crosslinked three times (6000 mJ, 245 nm), re-spreading between exposures.

### Heat shock

Mid-log cultures were rapidly shifted from 30°C to 40°C by brief incubation in a 70°C water bath with agitation, followed by incubation at 40°C for 30 min. Cells were harvested and frozen as described above.

### PRofiling Of Imprinted Transcripts (PROFIT) and subsequent purification of the Pol II-containing complex

3 liters of culture were grown to mid-log in YPD media at 30°C 200rpm. Pulverization, UV crosslinking, were performed as above. 7ml of cold lysis buffer was added to the grindate (50 mM Tris HCl, pH 7.4, with 150 mM NaCl, 1% TRITON X-100, 2.5 mM MnCl2) supplemented with 1X EDTA-free protease inhibitor cocktail (Roche). The lysate was centrifuged for 30 minutes,13,000rpm at 4°C. Chromatin associated proteins were extracted by adding 2ml lysis buffer supplemented with 2000 units of RNase free Dnase I (Sigma) to the pellet gently mixing. The lysate was incubated with the DNase for 1 hour on ice mixing periodically. The lysate was then centrifuged for 30 min, 13000rpm at 4°C. Supernatants were incubated on ice for an additional 30 minutes and then Dnase I digestion was stopped by 10 mM EDTA. Equal amount of protein were added to FLAG beads (Sigma). Lysates were incubated with the beads for 4 hours at 4°C with gentle agitation. The bead/lysate slurry was applied to a Biorad polyprep column. Beads were washed 4X with 1ml wash buffer (50 mM Tris HCl, pH 7.4, with 150 mM NaCl, 10mM EDTA) supplemented with Protease inhibitors Complete (Roche). Beads were incubated with wash buffer with 500 units RNase I (New England Biolabs) for 3 hours at room temperature with gentle agitation. The digest was eluted by centrifugation 2000rpm for 2 minutes at room temperature. Additional wash buffer was added to the beads to elute any residual proteins. Following the RNase I treatment, beads were washed 3X with 1ml wash buffer. Pol II associated proteins were eluted by excess FLAG peptide according to manufacturer’s instructions.

### Effect of *RPB4* deletion and mutations on PROFIT

For analysis of *rpb4Δ* effects, FLAG-tagged Rpb3 was purified from fragmented chromatin of WT and *rpb4Δ* strains. RNase-treated fractions were analyzed by LC–MS/MS. Label-free quantification (LFQ) data were log₂-transformed, proteins that did not appear in 2 (RNase-treated or Control) were filtered out, missing values imputed, and statistical significance assessed by t-test with permutation-based FDR correction (FDR = 0.05, S0 = 0.1) using Perseus.

### Mass Spectrometry

Trypsin-digested samples were analyzed by LC–MS/MS on a Q Exactive Plus (Thermo). Mass spectra were analyzed by MaxQuant with peak matching for the identification of proteins from different runs and compared with the theoretical yeast spectra produced from the Uniprot database and with a decoy reverse database. Statistical analysis was done using Perseus software n as follows: Label-free quant (LFQ) values MS analyses combining RNAse-treated and control were transformed to log2 scale. After filtering out proteins that did not appear in 3 repeats of at least one of the groups (RNase-treated or Control), missing values were imputed from normal distribution. The 2 groups were compared using t-test while applying permutation-FDR correction with FDR=0.05 and S0=0.1.

### Biotin labeling

Cells were grown to mid-log phase in biotin-free synthetic defined medium (1.7 g/L YNB-Biotin [Sunrise Science Products], 5 g/L Ammonium sulfate, 20 g/L dextrose, complete amino acids) supplemented with 0.125 ng/mL d-biotin at 24°C. For labeling, d-biotin was added to 1 µM for 15 min prior to harvesting, pulverization, and UV crosslinking.

### A small scaled and rapid detection of biotinylated proteins

Cells grown in biotin-free medium were induced with 1 µM biotin for the indicated times. 10 – 20 ml aliquots of 1×107 cells/ml were collected by centrifugation. Cell pellets were lysed in 50% urea, 7% SDS, 50 mM Tris-HCl pH 7.4, heated at 90°C for 10 min, and clarified by centrifugation. Supernatants were analyzed by western blotting.

### Western Blots

Proteins were separated on CriterionTM TGX Stain-FreeTM pre-cast polyacrylamide gel 4-15% (Bio-Rad Laboratories, California, USA) and transferred to PVDF membranes. To detect the bio-proteins, membranes were decorated with either 680nm infra-red (IR)-labeled streptavidin (IR-streptavidin) (IRDye® 680RD, Licor), to detect the bio-protein, or anti-AviTag Abs, using 800nm secondary labeled IR antibodies, to detect the bulk AviTagged protein. Membranes were scanned using LI-COR Odyssey M Imager at 700nm (for Streptavidin) and 800nm (for the remaining antibodies) wavelengths.

### BioPROFIT

BioPROFIT of biotin labeled cells (see above) was performed as PROFIT, with the addition of a desalting step (Zeba spin columns, Thermo) to remove free ATP and biotin prior to affinity purification.

### Polysome fractionation

Sucrose gradients (10–50%) were prepared and equilibrated overnight at 4°C. Yeast cultures were grown overnight either in 1L YPD or minimal Biotin media at 30°C until reaching cell density of 1*10^7^, pulse labeled with biotin as indicated in the figure legends, treated with cycloheximide (0.1 mg/ml), cryogenically pulverized (six cycles, 30 s, 15 Hz) using a Retsch MM301 mixer mill, and dissolved in lysis buffer (40 mM Tris HCl pH=7.4, 110 mM KCl, 1.5 mM MgCl_2_, 0.5 mM DTT, 0.8 mg/ml Heparin, 0.8% Triton X-100, 0.1 mg/ml cycloheximide and protease inhibitors), followed by clearance at 9000 g for 10 min at 4°C. One ml of 1 mg/ml protein was layered onto gradients. After ultracentrifugation (35,000g, 2.5 h, 4°C, using SW41 rotter), gradients were fractionated with UV detection (ISCO) at 254 nm. Proteins were precipitated with TCA and analyzed by SDS–PAGE and western blotting. Band intensities were quantified using Fiji.

## Supplemental figures

**Supplemental Figure 1.**
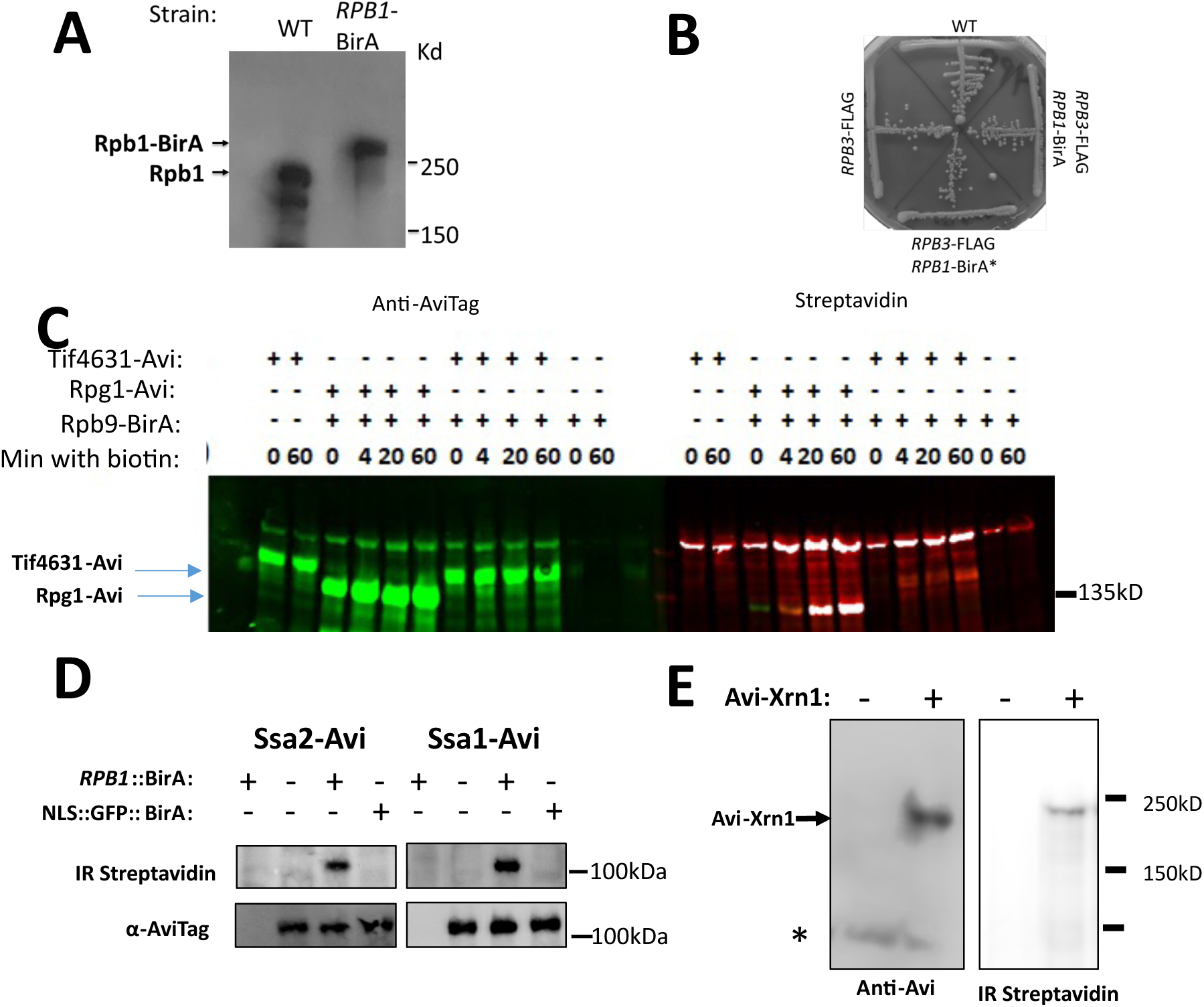
BirA fused to Pol II subunits specifically biotinylates AviTagged proteins. **(A-B) *Fusion of BirA to Rpb1 shifts electrophoretic mobility (A) without compromising cell growth (B)***. BirA was surgically inserted at the endogenous *RPB1* (A, B, D, E) or *RPB9* (C) loci, leaving both 5’ and 3’ non-coding regions unpurterbed. **(C) *Rpg1-AviTag and Tif4631-AviTag are biotinylated by Rpb9-BirA following biotin addition***. Cells were lysed under harsh denaturing conditions (including 50% urea, 10% SDS) to prevent post-lysis enzymatic activity and analyzed by western blotting. (D) ***Five minutes of biotin labeling is sufficient to detect AviTagged Ssa1 or Ssa2 in strains expressing Rpb1-BirA, but not NLS-GFP-BirA***. Bulk Ssa1 was detected with anti-AviTag antibody, while biotinylated Ssa1 (bioSsa1) was detected using IR-Streptavidin (see Materials and methods). **(E) *Xrn1-AviTag is biotinylated in vivo*.**

**Supplemental Figure 2.**
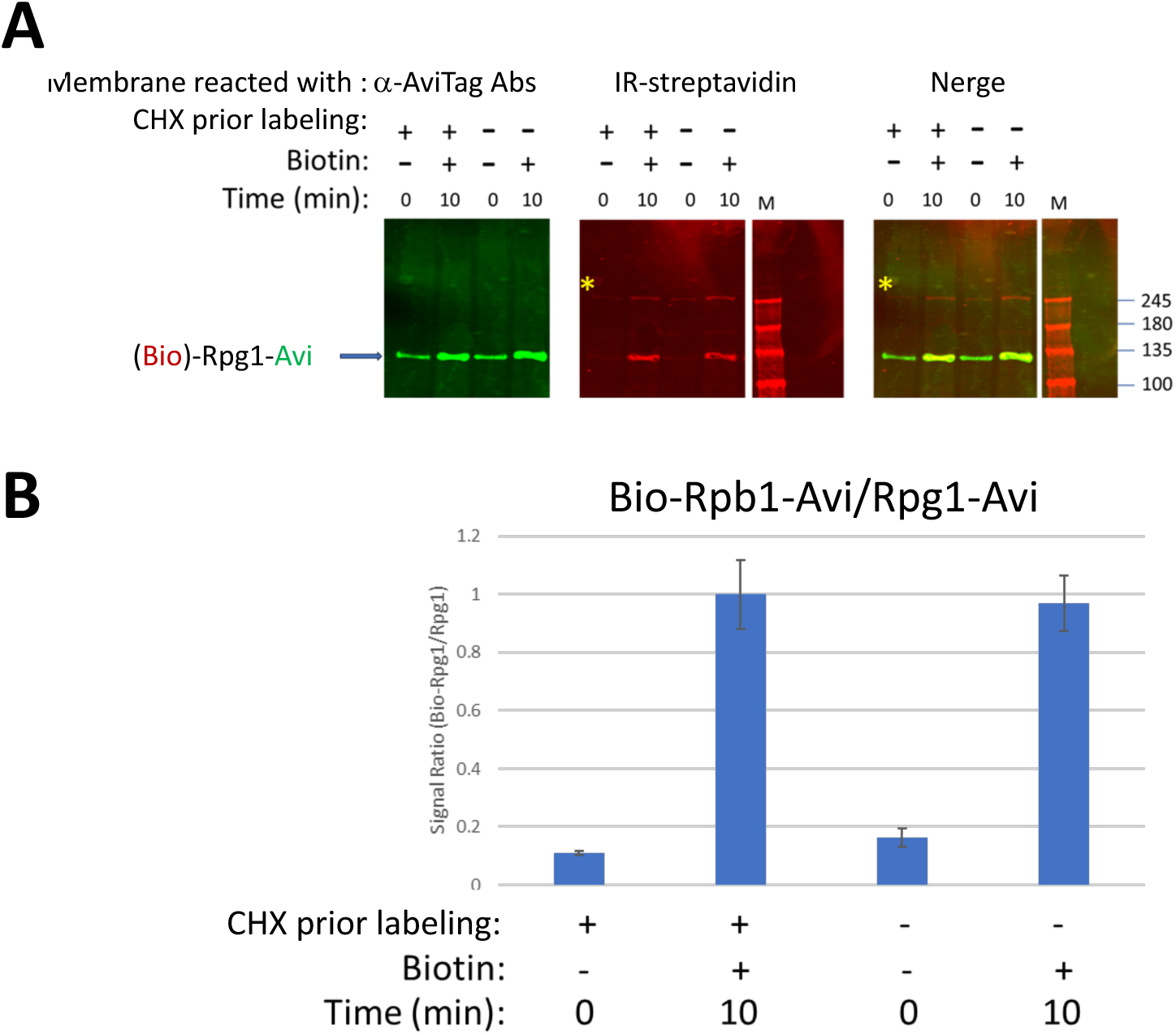
Translation inhibition does not alter Rpg1 biotinylation. **(A)** Rpb9-BirA Rpg1-AviTag cells were treated with cycloheximide (CHX) either before or after biotin labeling. Cells were lysed under denaturing conditions (50% Urea 10% SDS) to block enzymatic activity and analyzed by immunoblotting. **(B)** Band intensities were quantified (*Materials and Methods*), normalized to total signal, and used to calculate the ratio of biotinylated to total Rpg1. Data represent mean ± SD from two biological replicates.

## Notes

### Competing Interest Statement

The authors have declared no competing interest.

